# Molecular origins of transcriptional heterogeneity in diazotrophic *K. oxytoca*

**DOI:** 10.1101/2020.02.18.955476

**Authors:** Tufail Bashir, Rowan D Brackston, Christopher J Waite, Ioly Kotta-Loizou, Christoph Engl, Martin Buck, Jörg Schumacher

## Abstract

Phenotypic heterogeneity in clonal bacterial batch cultures is an important adaptive strategy to changing environments, including in diazotrophs with the unique capacity to convert di-nitrogen into bio-available ammonium. In diazotrophic *Klebsiella oxytoca* we simultaneously measured mRNA levels of key regulatory (*glnK-amtB, nifLA*) and structural (*nifHDK*) operons required for establishing nitrogen fixation, using dual molecule, single cell RNA-FISH. Through stochastic transcription models and mutual information analysis we revealed likely molecular origins for heterogeneity in nitrogenase expression. In wildtype and regulatory variant strains we inferred contributions from intrinsic and extrinsic noise, finding that *nifHDK* transcription is inherently bursty, but that noise propagation through signalling is also significant. The regulatory gene *glnK* had the highest discernible effect on *nifHDK* variance, while noise from factors outside of the regulatory pathway were negligible. Results provide evidence that heterogeneity is a fundamental property of this regulatory system, indicating potential constraints for engineering homogeneous nitrogenase expression.

## Introduction

Cell to cell variability in transcription has been recognised across many different cell types, and attributed to a range of causes (***Elowitz, 2002; Engl, 2019***). In bacteria such variability has been suggested to underpin phenotypic differences, or heterogeneity, between otherwise genetically identical cells cultured under a particular condition. Understanding this phenotypic heterogeneity in bacterial clonal populations has important implications for medical and biotechnological applications (***201, 2016***).

Phenotypic heterogeneity may be particularly relevant in costly stress response systems, of which the response to nitrogen starvation is a key example (***Schreiber et al., 2016***). In organisms such as *Klebsiella oxytoca*, nitrogen starvation triggers a transition to diazotrophic behaviour in which bacterial cells use atmospheric di-nitrogen as their nitrogen source for growth (***Dixon and Kahn, 2004***). While this transition, and the associated transcriptional programme, is essential for continued growth under conditions deplete of fixed nitrogen, it is also very costly since the resultant ATP consuming nitrogenase enzyme may ultimately constitute up to 20 % of the proteome (***Dixon and Kahn, 2004***). As such, if fixed nitrogen sources soon become available again, it is likely advantageous to have not fully undergone the diazotrophic transition. Activation of the nitrogen stress response is potentially somewhat of a gamble in which the payoff depends strongly on future conditions. Such a scenario may therefore indicate a requirement for bet-hedging strategies in which heterogeneity proves advantageous at the population level (***van Boxtel et al., 2017***).

Regardless of whether there are advantages to heterogeneity, it is also important to establish the underlying mechanistic causes, especially if there is an aim to ultimately modify cell behaviour. Studies at single-cell resolution are crucial to this, enabling the full distribution of expression levels to be obtained over a population of cells. Such data not only provides qualitative insight into the variability between cells, but can also enable the inference of the underlying dynamical processes (***Munsky et al., 2018***), through the combination of biological insight and quantitative models. This is the approach we take here to investigate the sources of noise that lead to heterogeneity of *nifHDK* gene expression at the transcriptional level.

In *K. oxytoca*, expression of a functional nitrogenase involves coordinated transcription of 18 *nif* genes that are organised in 5 operons. The *nifHDK* operon encodes the structural genes of the nitrogenase and is the most highly expressed operon within the *nif* cluster. The core hierarchical regulatory system of *nif* gene expression (Fig. 1(A)) consists of the nitrogen regulator NtrC activating expression of *glnK-amtB* and *nifLA* operons, with NifA activating *nifHDK* gene expression when not directly inhibited by NifL (***Dixon and Kahn, 2004***). Both NtrC and NifA are bacterial enhancer binding proteins (EBP) that activate the major variant σ^54^ RNA polymerase, with a distinct ATPase dependent activating mechanism compared with the canonical σ^70^ type RNA polymerases (***Schumacher et al., 2006***). The inhibitory NifL-NifA complex is destabilised by (i) GlnK binding, (ii) a reduced state of the NifL, (iii) ADP and (iv) α-ketoglutarate binding (reviewed in (***Dixon and Kahn, 2004***)). This arrangement integrates signals conducive to nitrogen fixation, namely a reducing environment, high energy levels and presumably low nitrogen levels, as high α-ketoglutarate levels indicate a low nitrogen status in the closely related *E. coli* (***Schumacher et al., 2013***). A low nitrogen status, defined as the ratio of glutamine/α-ketoglutarate also increases the activity of NtrC and enhances its expression by triggering uridylation of PII signalling proteins. Further, low glutamine levels affect the post-translational uridylylation state of GlnK, however it is unclear if the uridylation state affects GlnK function in destabilising the NifL-NifA complex in *K. oxytoca* (***He et al., 1998***).

**Figure 1.**
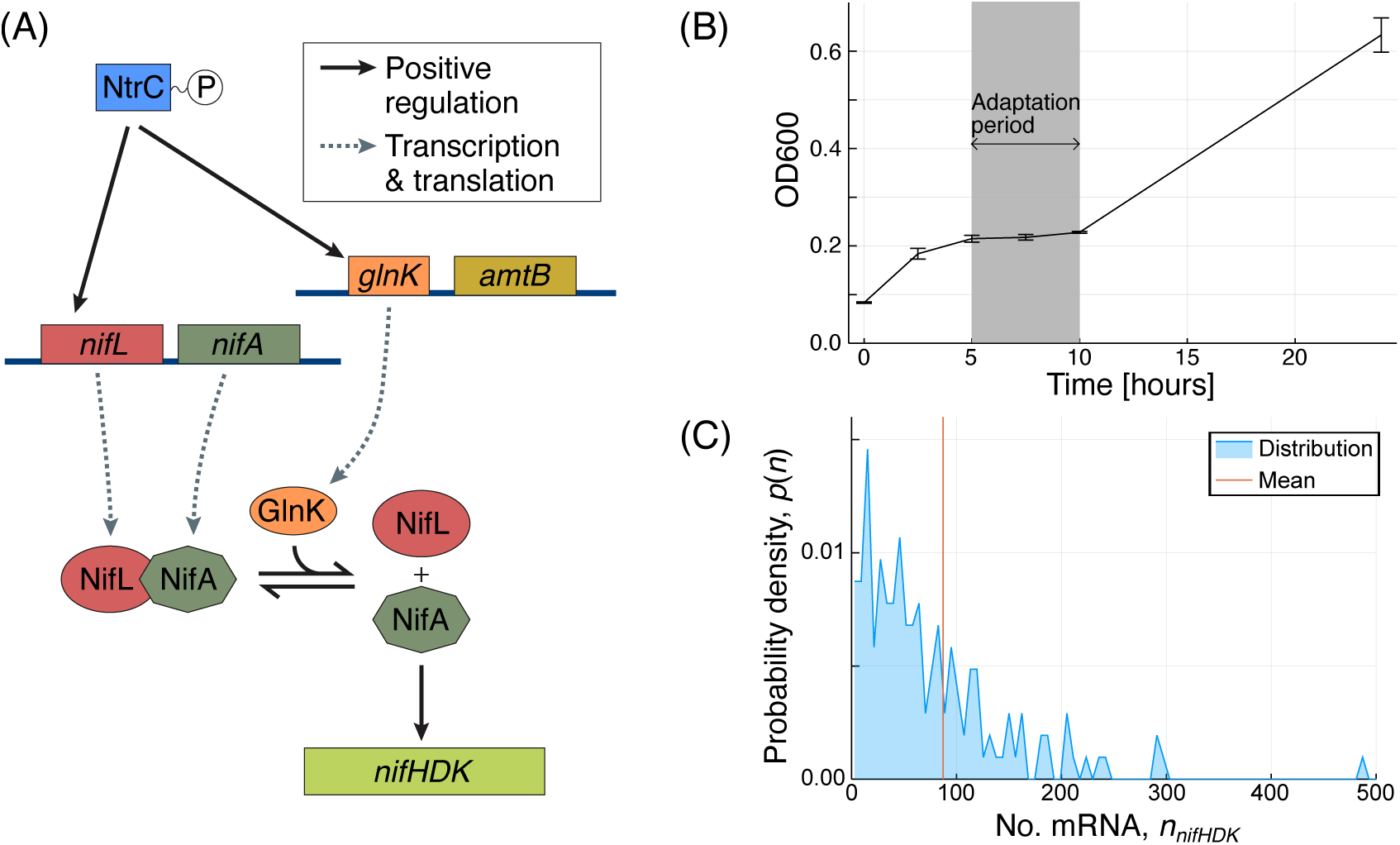
(A) Regulatory pathway governing *nifHDK* expression, adapted from ***Dixon and Kahn*** (***2004***). Transcription of *nifHDK* is subject to a hierarchical regulatory system. (B) Population size during transition to diazotrophy in wild-type *K. oxytoca*. Following run-out of ammonia, cultures display arrested growth during the diazotrophic transition, marked in gray. Growth from ten hours onwards is achieved through nitrogen fixation. (C) Distribution of *nifHDK* transcript abundance at 8 hours. Almost all cells have non-zero expression levels, but there is significant variability across the population. The average expression level is not necessarily representative of the typical cell. **Figure 1–Figure supplement 1.** Cumulative acetylene reduction comparison.

Phenotypic heterogeneity in nitrogen fixing *K. oxytoca* grown in chemostats has been demonstrated under limiting ammonium availability and suggested to be caused by events downstream of GlnK (***Schreiber et al., 2016***). A degree of phenotypic heterogeneity is known to result from the inherent stochasticity in gene expression and has been widely observed in other systems (***Elowitz, 2002; Cai et al., 2006; Kiviet et al., 2014***). Such stochasticity is common to all chemical reaction systems involving small numbers of molecules of which transcription is a key example. However transcription is further observed to occur in bursts (***Jones and Elf, 2018; Golding et al., 2005; Suter et al., 2011; Larson et al., 2013***), short periods of intense transcriptional activity, resulting in in-creased levels of heterogeneity. Together, inherent stochasticity and burstiness lead to what is often referred to as intrinsic noise which may be a fundamental property of transcription of a given gene. However, it is understood that in addition to this intrinsic noise, other sources of noise external to a particular gene may also be relevant. These additional contributions to heterogeneity, referred to as extrinsic noise, have been observed experimentally by simultaneously measuring the expression of two or more copies of the same gene at the single cell level (***Swain et al., 2002; Raser and O’Shea, 2004; Gasch et al., 2017***). Correlations in the expression of these two gene copies reflect perturbations that simultaneously affect both.

While noise contributions can sometimes be directly measured, it is also possible to make inferences via a modelling approach. Intrinsic noise and bursty transcription have long been modelled by the so-called Telegraph process that describes stochastic events within transcription and predicts the probability distribution of mRNA copy numbers (***Ko, 1991; Peccoud and Ycart, 1995; Raj et al., 2006; Iyer-Biswas et al., 2009***). This model and its variants have been widely used to infer transcriptional properties (***Schreiber et al., 2016; Munsky et al., 2015; Sepúlveda et al., 2016; Larsson et al., 2019***) and therefore provide improved analysis and understanding from experimental data. Such models can also be readily extended to include the effect of extrinsic noise (***Lenive et al., 2016; Sherman et al., 2015; Ham et al., 2019***), and subsequently quantify this contribution.

Here we are interested in the relative contributions of intrinsic and extrinsic noise to expression of the *nifHDK* operon, under conditions in which free living *K. oxytoca* cells transition to use atmospheric di-nitrogen as their nitrogen source for growth. Using smRNA-FISH and examining the relationship between genes at different levels in the regulatory cascade controlling nitrogenase gene expression, we are able to reveal the combined roles of intrinsic noise at the level of the *nifHDK* promotor and extrinsic noise arising from upstream regulation. By further fitting stochastic models for the transcription process, these contributions are quantified, along with additional details of the regulatory mechanisms.

## Results

### Transition to diazotrophy

To measure the full spectrum of heterogeneity pertaining to the transition into and establishment of full diazotrophy, precultures grown in nitrogen replete aerobic conditions were transferred to nitrogen free, anaerobic media at time zero, when no *nif* expression or acetylene reduction was detectable. We found that *K. oxytoca* (WT) populations subjected to these conditions first consumed residual ammonia then experienced a growth arrest, which lasted around 5 hours, (see Fig. 1(B)). This was followed by a resumption of growth using N_2_ gas derived ammonia as their nitrogen source, evidenced using the established acetylene reduction activity profiles as a measure of bulk population nitrogenase enzyme activity in batch culture (***Dilworth, 1966***).

In order to investigate and verify the regulatory roles of *glnB, glnK* and *nifA*, we also grew cells lacking the positive regulator GlnK (Δ*glnK*), cells lacking the enhancer binding protein NifA and its co-transcribed inhibitor NifL (Δ*nifLA*), and cells lacking the upstream regulator GlnB (Δ*glnB*), which controls the NtrB dependent phosphorylation of NtrC in the cells in diazotrophic behaviour. We conducted the same experiments on each mutant, tracking the population growth and Nitrogenase activity (see Fig. 1 S1). We found that the absence of GlnB had negligible effect. By contrast both the Δ*glnK* and Δ*nifLA* mutants showed markedly different behaviour to the wild-type, also confirming the known role of each. Some nitrogen fixation was evidenced in the absence of GlnK (30 % of that in WT at 9.5 hours), but the Δ*nifLA* mutant was unable to fix at any level.

Further to these population averaged observations, we established use of FISH in *K. oxytoca* to estimate single molecule mRNA levels for the *nifHDK* operon which encodes the iron protein complex of the nitrogenase enzyme. We followed the protocol outlined in (***Skinner et al., 2013***), measuring expression in two additional control samples, as detailed in materials and methods. Results indicated significant variation in *nifHDK* transcript abundance (see Fig. 1(C) for an example). As is common in gene expression data, the distribution is decidedly non-Gaussian, with a bulk of cells at low expression levels and a longer tail of much higher values. This means that the “average” cell has relatively high expression levels compared with the majority, and therefore that bulk measurements of *nifHDK* gene expression do not represent a typical cell.

### Mutual information analysis confirms direct propagation of transcriptional noise to *nifHDK* via GlnK

Given the observation of significant heterogeneity in *nifHDK* gene expression, we sought to establish whether extrinsic noise is present, and if this noise arises from the regulatory pathway or from elsewhere. We therefore grew WT cells, Δ*glnK* and Δ*glnB* mutants, to levels where different bacterial cultures displayed equivalent cumulative levels of nitrogenase activity (as described in methods). Two sets of probes were used to measure *nifHDK* and simultaneously either *glnK* or *nifLA* expression in individual cells to monitor variability across these paired genes (see Fig. 2 S1). Generally speaking, expression co-variance between two genes can either occur because one gene has a direct (and relatively quick) influence on the other gene, or because both genes are simultaneously affected by another source of variability. This could be a shared upstream control protein, or “global” factors such as RNA polymerase or sigma factor abundance that may influence many genes simultaneously and should therefore be evident as significant co-variance between any expressed pair of genes.

**Figure 2.**
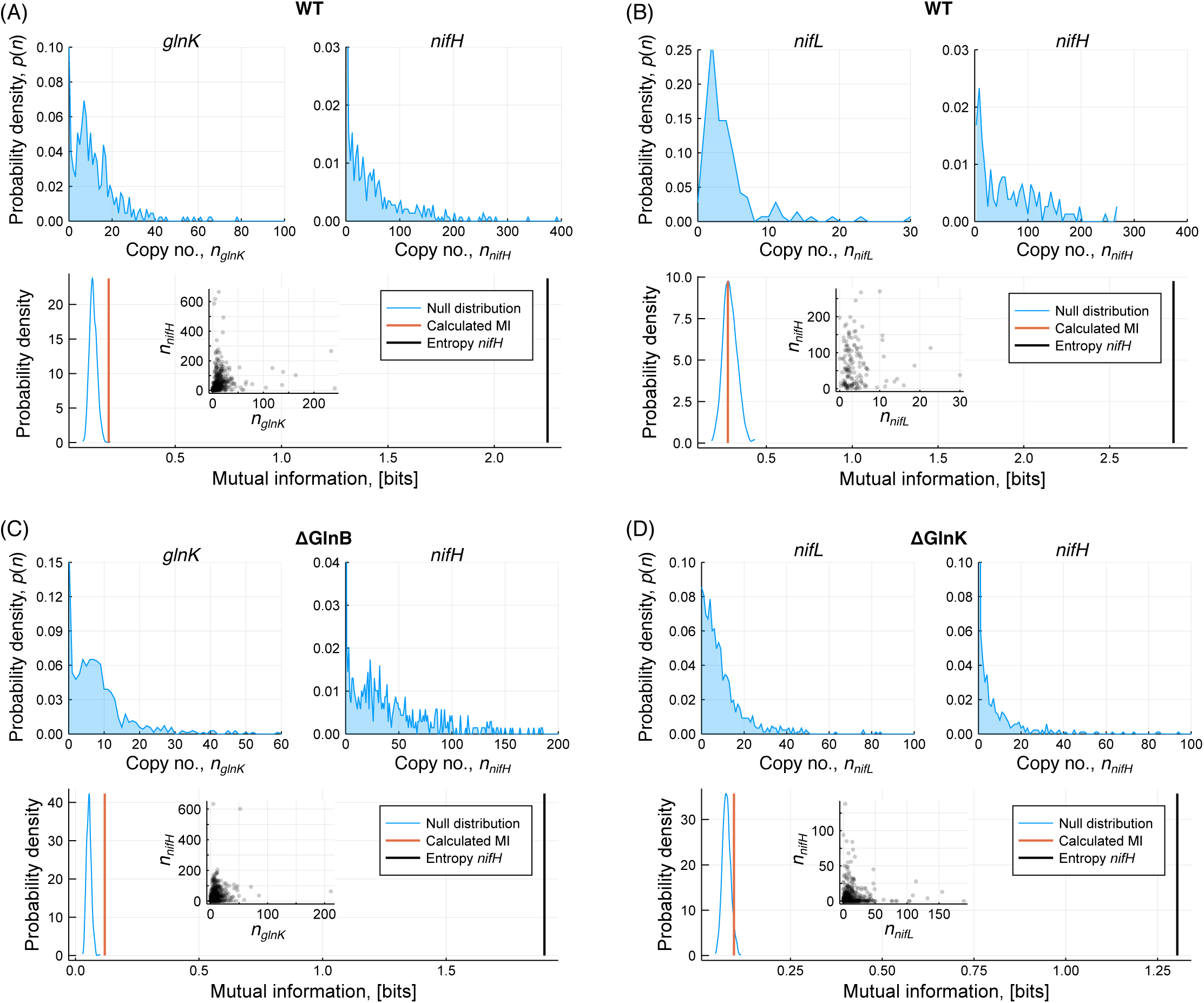
Mutual information analysis based on dual probe measurements for one set of biological replicates. See supplementary figure for exemplar images. For each case the marginal distributions of the two mRNA abundances are shown, in addition to the calculated mutual information and the total entropy in *nifHDK* abundance. Mutual information is compared to a null distribution obtained by randomly shuffling the *nifHDK* data 100,000 times, thereby providing a *p*-value for each pair. These values for the displayed data and a further biological replicate are displayed in Tab. 1. **Figure 2–Figure supplement 1.** Raw images demonstrating the use of dual FISH probes.

We used mutual information (MI) as a suitable metric for quantifying co-variance in gene expression. MI is commonly used to analyse single cell data in situations where relationships are complex, nonlinear and unknown, as no assumptions or prior knowledge are required (***Mc Mahon et al., 2014, 2015***). Here, we compute the MI between each pair of genes and a null distribution of MI values from which a corresponding significance level can be computed (see materials and methods). The MI can further be compared to the entropy of the distribution of *nifHDK* expression levels, in order to compare the mutual information with the total information content. The results of this analysis are displayed in Fig. 2 and Tab. 1.

**Table 1.** Mutual information with *nifHDK* transcript abundance. The entropy of *nifHDK* abundance gives a measure of the total variability, while the mutual information is the amount of variability that can be directly explained by the abundance of the other gene. Smaller significance levels indicate a stronger statistical indication of non-zero mutual information.

| Genetic strain | Gene | Mutual information [bits] | Entropy *nifHDK* [bits] | Significance level |
| --- | --- | --- | --- | --- |
| WT | *glnK* | 0.188 | 2.25 | 0.00016 |
| WT | *glnK* | 0.340 | 2.44 | 0.00035 |
| WT | *nifL* | 0.275 | 2.87 | 0.62 |
| WT | *nifL* | 0.318 | 2.75 | 0.47 |
| $\Delta glnB$ | *glnK* | 0.118 | 1.90 | 0.0 |
| $\Delta glnB$ | *glnK* | 0.072 | 1.92 | 0.0074 |
| $\Delta glnK$ | *nifL* | 0.067 | 0.69 | 0.28 |
| $\Delta glnK$ | *nifL* | 0.096 | 1.30 | 0.047 |

The results demonstrate a modest level of MI between *glnK* and *nifHDK* but no statistically significant measure of MI between *nifLA* and *nifHDK*. Undetectable MI between *nifLA* and *nifHDK* suggests that global sources of extrinsic noise are not significant for these two genes, and by extension are unlikely to be significant for any other σ^54^ dependent genes. The measurable low-level MI between *glnK* and *nifHDK* is therefore suggestive of a direct propagation of noise from *glnK* to *nifHDK*. In other words, stochastic variability of *glnK* expression acts to increase the level of variability in *nifHDK* expression; those cells that have high levels of *glnK* mRNA at a given time, are also likely to have higher levels of *nifHDK* mRNA. While this is clearly intuitively consistent given the known direct role of *glnK* to de-repress the inactive NifL-NifA complex (***He et al., 1998; Little et al., 2002***), the implications for a lower bound on *nifHDK* variability are significant. Given that the cells are exposed to identical conditions within the culture, it is noteworthy that variability at the level of *glnK* acts to increase variability in *nifHDK*.

### Stochastic models incorporating extrinsic noise

Next, we sought to understand the minimum level of variability that might be observed if the direct regulatory role of GlnK were bypassed. This was done by measuring expression levels in a Δ*nifLA* strain in which *nifA* is overexpressed ectopically on a plasmid (+*nifA*, see materials and methods), thus making *nifHDK* expression independent of GlnK control.

Fig. 3(A) displays a distribution for the +*nifA* mutant, showing significant remaining sources of heterogeneity. Given that fluctuations in GlnK are no longer anticipated to have an effect on *nifHDK* expression, and that global sources of extrinsic noise were found to be minimal, *nifHDK* transcription heterogeneity must be due to intrinsic variability at the level of the *nifHDK* promotor. Such variability is widely attributed to bursty transcription, in which the promotor is intermittently active and inactive (***Jones and Elf, 2018; Golding et al., 2005***). The sources of this intermittent activity may in turn be related to transcription factor binding/unbinding (***Jones et al., 2014***), or to mechanical supercoiling effects (***Chong et al., 2014; Sevier et al., 2016***). Regardless of the mechanism, it is likely that similar levels of intrinsic variability may also be relevant when other genetic variants are tested.

**Figure 3.**
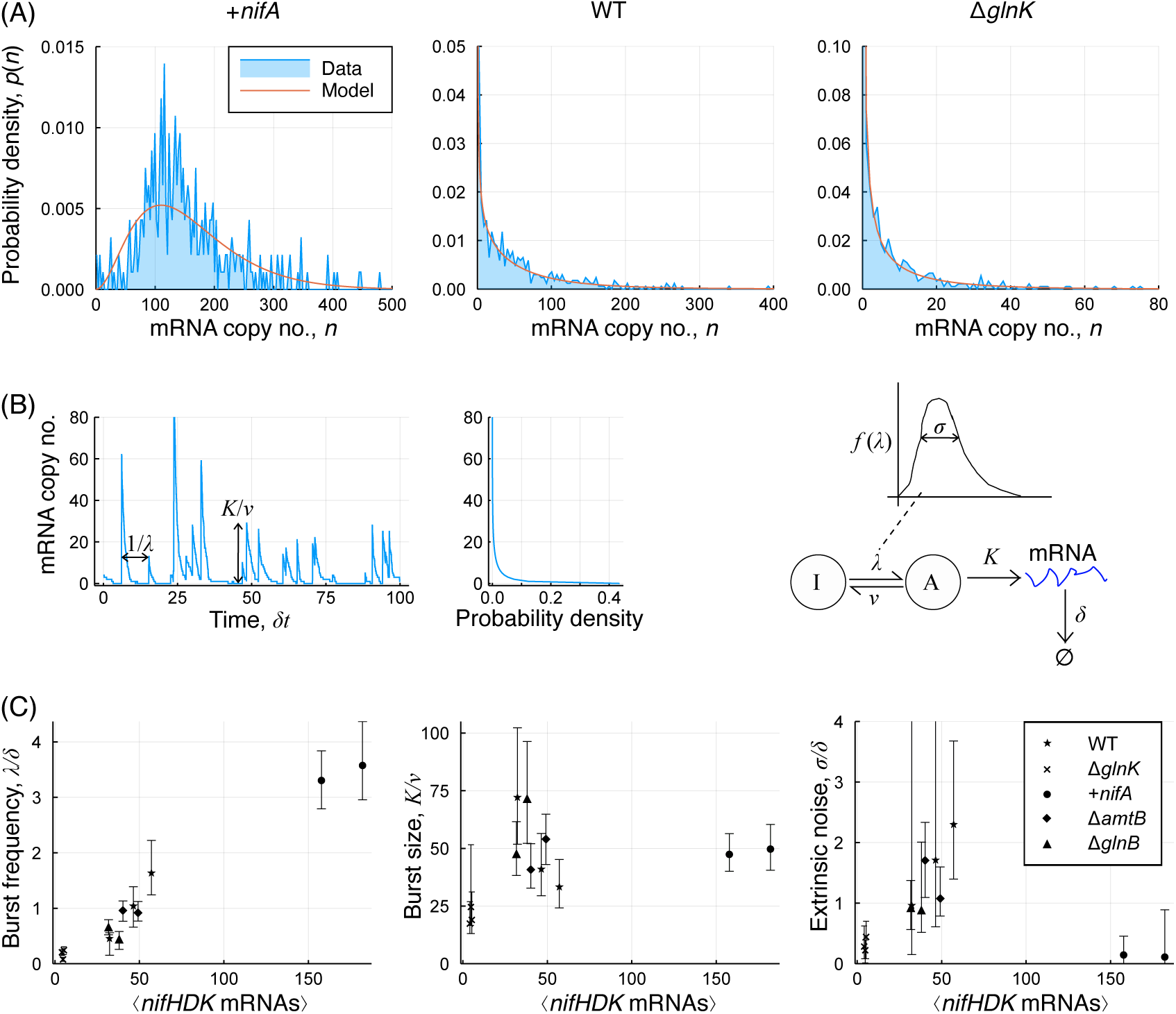
Stochastic modelling of bursty transcription incorporating extrinsic noise. (A) mRNA copy number distributions for each of the +*nifA*, WT and Δ*glnK* strains. (B) Schematic of bursty transcription and its relation to the stochastic model. Extrinsic noise here is catered for by considering the burst frequency to itself be variable between cells. (C) Variation of the model parameters between mutants, plotted as a function of the mean expression level. Error bars denote 95% Bayesian credible intervals, obtained from the MCMC chains (see figure supplement.) **Figure 3–Figure supplement 1.** Exemplar MCMC chain and posterior distributions from parameter inference.

Based on the observations that both intrinsic and extrinsic sources of noise may be generally relevant, we sought to develop stochastic models for transcription that could incorporate both these effects, following the approach outlined in (***Ham et al., 2019***). When considering *nifHDK* transcription to be intrinsically bursty, as shown schematically in Fig. 3(B), this can be modelled by the Telegraph model (***Peccoud and Ycart, 1995; Iyer-Biswas et al., 2009***), where the promotor undergoes rapid activation and deactivation at rates *λ* and *ν* respectively. When active, transcription occurs at a rate *K* while the mRNA is degraded according to a first order degradation process with rate *δ*. It can be shown that under particular parameter ranges this process results in a negative binomial distribution for the transcript copy number (***Ham et al., 2019; Paulsson and Ehrenberg, 2000***), parametrized by the normalised burst frequency *λ*/*δ* and mean burst size *K*/*ν*. In this context we take the view that extrinsic noise arising from *glnK* variability acts as a variation between cells in the burst frequency. This leads to a model that is additionally parametrized by the normalised frequency variation *σ*/*δ*. Further details are given in the materials and methods.

For each of the three exemplar distributions shown in Fig. 3(A), a comparison is given with the model following the parameter fitting process. It is evident that the model can provide a good fit to the data in each case, thereby enabling us to draw meaning from the model parameters, inferred for a number of genetic variants and displayed in Fig. 3(C). We observed a clear relationship between the mean expression level and the burst frequency, which is contrasted by the very limited correlation between mean expression and burst size. Cells with very low expression levels such as the Δ*glnK* strain have a very low burst frequency, yet their burst size is within a factor of two of that for the most highly expressing +*nifA* strain. For the lower expressing strains, the level of inferred variability in the burst frequency seems proportional to the average. The same is not true for the +*nifA* strain in which extrinsic noise is very low.

Based on these model fits, we further calculated the contributions of intrinsic and extrinsic noise to the total variance in *nifHDK* expression levels (***Sherman et al., 2015; Hilfinger and Paulsson, 2011***) (see methods), as shown in Fig. 4(A). For the Δ*glnK* strains, the results demonstrate that there remains a significant contribution from extrinsic noise of around 30%, indicating that some variability in burst frequency may arise from noise sources other than cell to cell variation on GlnK activity. The extrinsic noise contribution increases to around 70% in the WT, Δ*amtB* and Δ*glnB* mutants in which noise propagation from *glnK* is known to be present, but is almost zero in the +*nifA* mutant in which the regulatory pathway via GlnK is uncoupled.

**Figure 4.**
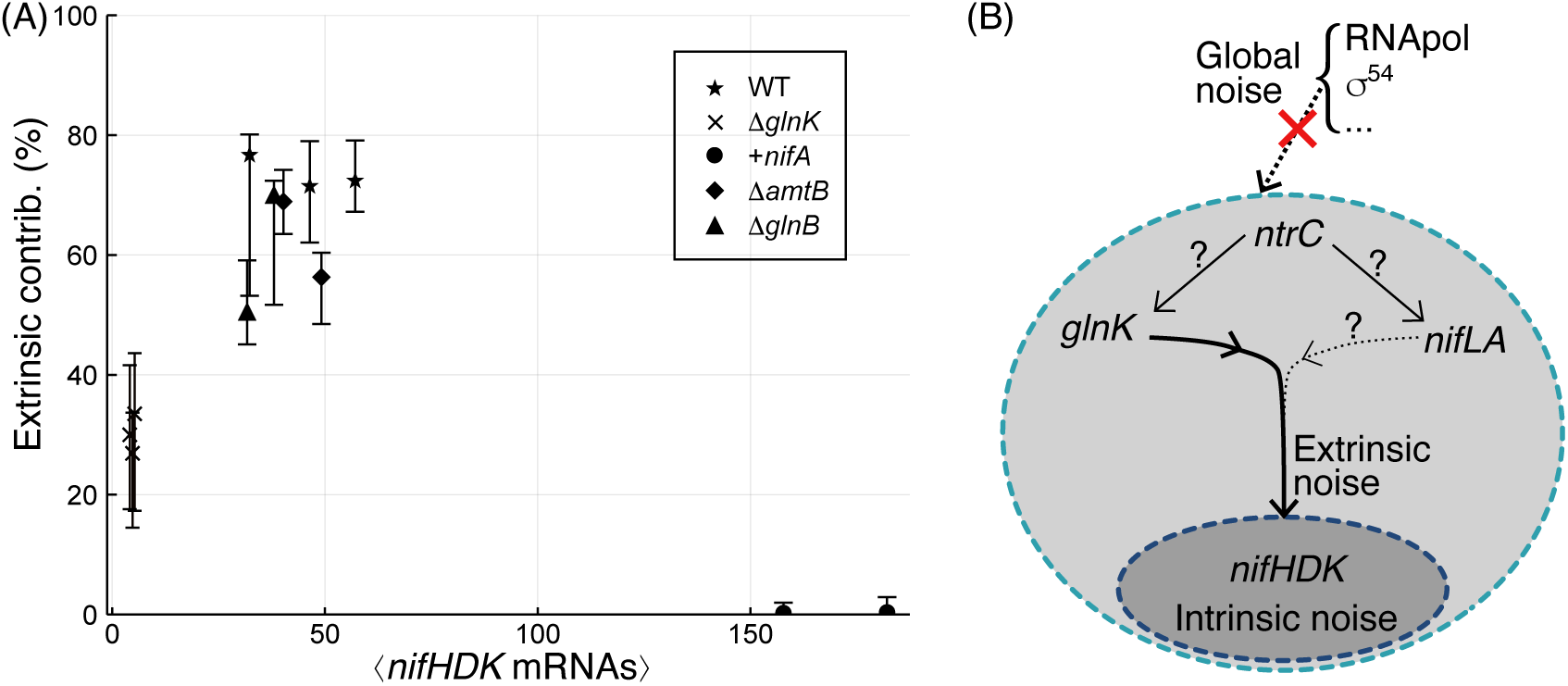
(A) Calculated contributions of extrinsic noise to the variance in *nifHDK* transcript abundance. Contributions are calculated for thousands of parameters sampled from the MCMC chains, thereby providing the most probable value and 68% credible intervals. (B) Schematic showing propagation of noise through the signalling cascade. Only a contribution from *glnK* is supported directly by the experimental data, although contributions from *nifLA* and from *ntrC* cannot be excluded. Global noise sources that are found to be very small.

## Discussion

Bacteria can be subject to any of a multitude of stresses and respond in a multitude of ways. While the average behaviour of a clonal population is often of interest, the degree of variability between cells is also important. Such knowledge may provide both mechanistic biochemical insight, as well as an indication of typical survival strategies that leverage this variability. Here we have examined the nitrogen starvation response in a model diazotroph, determining behaviour at the population level and the expression of several genes in the regulatory cascade at single cell resolution.

From the bulk measurements, our data extends the existing understanding of the *nif* regulatory network in *K. oxytoca*. We find that a Δ*glnB* strain behaves in a similar manner to the WT in a number of respects, consistent with the highly homologous GlnK being able to compensate for the loss of GlnB. GlnB and GlnK were shown to have redundant functionalities in regulating nitrogen assimilation genes in *E. coli*, but this redundancy is not fully extended to *nifHDK* gene regulation in the closely related *Klebsiella pneumoniae* (***Arcondéguy et al., 1999***). Further, GlnB and GlnK post translational uridylylation plays an important role in regulating nitrogen assimilation genes but GlnK uridylylation is not required for *nifHDK* gene regulation (***He et al., 1998***). Our Δ*glnK* strain showing much slower *nifHDK* expression is consistent with these previous findings. Given that GlnK is largely required to dissociate NifA from the co-transcribed NifL, non-zero expression of *nifHDK* in Δ*glnK* must be indicative either of low-level spontaneous dissociation of NifA or compensatory action by GlnB. In *K. pneumoniae*, GlnB when overexpressed can functionally compensate for a deletion of *glnK* (***Arcondéguy et al., 1999***), and because *glnB*, unlike *glnK*, is constitutively expressed, such a compensatory mechanism of *glnK* by *glnB* could explain the reduced contribution of extrinsic noise. Lastly, low-level *nifHDK* expression did not occur in the Δ*nifLA* strain. This is consistent with a scheme of control in which NifA provides the essential but leaky switch for *nifHDK* expression while GlnK is the regulatory modulator.

By examining Mutual Information between pairs of genes we can infer how noise is propagated through the regulatory cascade. MI is generally small compared with *nifHDK* entropy and is below the level of statistical significance between *nifLA* and *nifHDK*, suggesting that global sources of extrinsic noise are negligible. In other gene transcription systems it has been suggested that variability in sigma factor or RNA-polymerase abundance can provide a source of variability in transcription (***Elowitz, 2002; Swain et al., 2002; Taniguchi et al., 2010***), but our results suggest that this is not the case here. We cannot exclude the possibility that fluctuations do propagate from NifLA, since the differences in mRNA and protein lifetimes can act to reduce correlations between transcript and protein abundance (***Taniguchi et al., 2010***). Fluctuations could also propagate from the master regulator NtrC, but any sources of noise above this level must be small and therefore all variability in *nifHDK* expression must arise from within, and not outside, the regulatory network (see Fig. 4(B)).

Unlike for *nifLA*, small but statistically significant MI is observed between *glnK* and *nifHDK*. This observation, together with the known positive regulatory role of GlnK, indicates that variability in *glnK* expression must propagate down to *nifHDK*. We therefore have a direct detection of extrinsic noise acting on *nifHDK* transcription.

While the MI analysis evidences the non-zero contribution of extrinsic noise to heterogeneity in *nifHDK* transcription, we further quantify this contribution through stochastic modelling. All single cell measurements for *nifHDK* expression are consistent with a model in which transcription is inherently bursty, even in the +*nifA* mutant in which the promotor should be constitutively active. This burstiness may result for example from unavoidable dissociation of the NifA EBP and acts to generate significant levels of intrinsic noise. The model also incorporates propagation of noise down the signalling cascade as variability in the frequency of the transcriptional bursts. Further to this, by examining the variation of model parameters across mutants we can deduce that the average expression level is principally determined by the burst frequency rather than the burst size. This is consistent with recent observations for the phage shock protein (Psp) membrane stress response which is also σ^54^ dependent (***Engl et al., 2019***), and in contrast to the existing understanding for σ^70^ promotors (***So et al., 2011***).

The model and inferred parameter variation displayed in Fig. 3(C) are consistent with a picture in which bursts are initiated by binding of the NifA EBP to its target promoter closed complexes and terminated by its unbinding. Extrinsic noise then relates to variability in the availability of the free NifA protein. In the Δ*glnK* strain most NifA proteins are bound to NifL meaning that bursts are rare, yet when they do occur, the duration of the burst is relatively unaffected. The absolute magnitude of the extrinsic noise in this case is small, since all cells are subject to similarly low NifA availability. For the opposite case of the +*nifA* mutant, high NifA availability enables frequent bursts of transcription, yet the burst size is similar to the WT and other mutants, perhaps because the average time before dissociation of NifA from the promotor is independent of its abundance within the cell. This may reflect the rather slow conversion of a closed promoter complex as bound by NifA to an open promoter complex from which NifA has dissociated (***Friedman et al., 2013***). The level of extrinsic noise in this case is very low, since NifA availability is large enough for the system to essentially be “saturated”: any variability in NifA has little impact if the burst frequency is already at a maximal value.

The modelling and quantitative analysis gives us mechanistic insight into the sources of heterogeneity, implying that the large variation in *nifHDK* expression is an inherent property of this system. What is not clear is the implication of this for evolutionary fitness. A first possibility is that the heterogeneity is either unavoidable or sufficiently benign that the extra regulatory effort that would be required to suppress it is not worthwhile (***Lestas et al., 2010; Yan et al., 2019***). In this scenario heterogeneity is no more than an interesting artefact of the regulatory system. The alternative is that heterogeneity is beneficial, perhaps in a bet-hedging sense, as has been indicated before in bacterial stress response systems (***Carey et al., 2018; Patange et al., 2018***). Actually testing for evidence of bet-hedging is generally challenging (***Simons, 2011; Grimbergen et al., 2015***), and the strength as a strategy depends both on the specifics of the stress response system and on the typical frequency of environmental changes (***Kussell and Liebler, 2005***). Since the response to nitrogen starvation incurs such a heavy metabolic cost and resource availability for enteric bacteria is highly unpredictable, bet-hedging remains plausible.

It is quite possible that the observations made here are applicable to many other stress response systems. However particular interest in the nitrogen starvation response arises from the desire to engineer higher bulk levels of nitrogen fixation (***Gasperotti et al., 2020***). In this context the significant heterogeneity observed here may impose particular limitations on industrial scale use of diazotrophs (***Delvigne et al., 2014***), as well as confound the efficient use of clonal populations of diazotrophs in the rhizosphere unless engineered to avoid variance. Our results suggest that heterogeneity can be somewhat reduced by bypassing *glnK* and expressing *nifA* heterologously, yet only to a degree. Further questions then arise as to the extent this would affect population level fitness. If the sustainable population size or geometric growth rate are affected too much, a homogenously fixing population may perform no better in bulk. We hope therefore that the results presented here will motivate a more careful consideration of the challenges surrounding the use of diazotrophic bacteria as a source of fixed nitrogen.

## Materials and methods

### Bacterial Strains and growth conditions

All experiments were performed with Klebsiella oxytoca M5aI, obtained from (***Yu et al., 2018***). Whole gene knockout mutants, marked with a kanamycin resistance (nptII) gene, were derived from M5a1 using Lambda red recombineering (Datsenko Wanner). To generate a strain overexpressing NifA, the M5aI *nifA* gene sequence was cloned into the pSEVA424 vector (seva.cnb.csic.es) under the control of the Ptrc promoter and a synthetic ribosome binding site (BBa_B0032, Registry of Standard Biological Parts), prior to transformation into the Δ*nifLA* mutant background by electroporation. NH_4_ run-out was used to de-repress *gln*/*nif* gene expression and stimulate a reproducible transition into diazotrophic growth. Briefly, *K. oxytoca* strains were cultured in Nitrogen-Free David and Mingioli (NFDM) medium (***Cannon et al., 1974***) [69 mM K_2_HPO_4_, 25 mM KH_2_PO_4_, 0.1 mM Na_2_MoO_4_, 90 μM FeSO_4_, 0.8 mM MgSO_4_, 2 % w/v glucose] supplemented with NH_4_Cl as a nitrogen source. To ensure replete cellular N status, seed cultures were supplemented with 20 mM NH_4_Cl and grown to an OD_600_ of approximately 2-3. Cells were washed and resuspended in NFDM supplemented with 0.5 mM NH_4_Cl to an OD_600_ of 0.1. Cultures were crimp-sealed in 70 ml glass serum bottles (Wheaton) and chilled on ice whilst sparged with N_2_ gas for 45 minutes to establish a micro-aerobic atmosphere. Colorimetric O2xyDot sensors (OxySense) fixed inside the bottles were used to verify O_2_ concentration. Following injection of 1 mL pure, O_2_-free acetylene into the headspace, cultures were warmed to 25°C and shaken at 200 rpm for up to 24 hours.

### Nitrogenase assay

Nitrogen fixation was assessed via the acetylene reduction assay (***Shah and Brill, 1973***): 500 μl of culture headspace was sampled via gas-tight syringe and subject to gas chromatography through a HayeSep N column (Agilent) at 90°C in N_2_ carrier gas. Acetylene and ethylene were detected by flame ionisation (FID) at 300°C and ChemStation software (Agilent) was used to integrate signal peak areas. Periodically, 15 ml of oxygen-free N_2_ gas was injected into sample bottles via gas-tight syringe prior to extraction of an equivalent volume of cell culture for analysis of OD_600_ and smRNA-FISH. Accumulative nitrogenase activity are expressed as % acetylene consumption and ethylene production, normalised by OD_600_.

### RNA fluorescence in-situ hybridisation

mRNA-FISH was performed according to the protocol described by ***Skinner et al.*** (***2013***). Briefly, bacterial cells were sampled anaerobically and collected by centrifugation. Pelleted cells were fixed in a buffer containing 3.7 % (v/v) formaldehyde prior to permeabilization in 70 % (v/v) ethanol. Hybridization and wash steps were performed in saline and sodium citrate buffer (SSC; 150 mM sodium chloride, 15 mM sodium citrate) supplemented with 40 % (v/v) formamide. All solutions were prepared using DEPC-treated water and RNase-free plasticware. DNA probes against the *nifHDK, nifLA* and *glnKamtB* structural operons were designed using the Stellaris® Probe Designer version 4.2; the oligo length was set at 20 nt, the minimal spacing length at 2 nt and the masking level at 1-2. The probes were purchased pre-labelled with 6-carboxytetramethylrhodamine, succinimidyl ester [6-TAMRA] (*nifLA, glnKamtB*) or Cy5 equivalent Quasar® 670 (*nifHDK*) from LGC Biosearch Technology. Hybridization was performed overnight at 30°C at a final concentration of 1 μM, in buffer containing 2 mM ribonucleoside-vanadyl complex (VRC), 1 mgml^-1^ *E. coli* tRNA and 10 % (w/v) dextran sulphate. Following multiple wash steps, chromosomal DNA was stained with 10 μgml^-1^ DAPI (4’,6-diamidino-2-phenylindole) for 30 minutes before cells were immobilised using 1 % (w/v) agarose pads on 35 mm high μ-Dishes (ibidi) for imaging.

### Microscopy and image analysis

Cells were harvested for carrying out RNA FISH and microscopy analysis as described in ***Skinner et al.*** (***2013***), with slight modifications e.g. 35 mm IBIDI discs were used for imaging purpose with the help of a WF1 Zeiss Axio observer inverted microscope. Multiple fields of view were acquired for each sample, and within each field of view five z-slices were captured for further processing. Data stacks were converted to TIFF format using ImageJ, and cell segmentation masks from brightfield or DAPI images were generated using Schnitzcells. The protocol outlined in ***Skinner et al.*** (***2013***) was then followed to detect and quantify mRNA in each cell using the Spatzcells package in MATLAB. Output from this software consisted of an estimated number of mRNA in each imaged cell.

We harvested cells from different bacterial strains when they exhibited the same cumulative levels of nitrogenase activity corresponding to 5 % acetylene reduction. Time points selected to achieve this level of nitrogenase activity across different M5a1 bacterial strains were 14.5 h for Δ*glnB* and M5a1 (WT) strain, 19.5 h for Δ*glnK* strain and 14.5 h for Δ*amtB* strain.

### Mutual information analysis

In order to assess the relationship between transcript abundance of two different genes, we evaluated the mutual information (MI). MI provides a method for evaluating statistical dependencies between two random variables from the joint and marginal probability densities, even when the underlying relationship is complex and nonlinear.

As with any statistic intended for classification, it is important to ascertain a confidence threshold in order to rule out false positives. This is achieved by evaluating a null distribution for the value of the test statistic in the case that there is no relationship. Here we achieve this by shuffling the data for one gene and evaluating the MI obtained in this case. By performing this shuffling and evaluation 100,000 times, a null distribution for the MI is obtained. From this null distribution a significance level can be obtained for the measured MI.

### Stochastic modelling and parameter inference

The stochastic model for transcription is based upon the Telegraph model in which a given gene transitions from inactive to active at rate *λ* and from active to inactive at rate *ν*. When the promotor is active, transcription occurs at rate *K*, while degradation of the mRNA occurs at rate *δ* independent of the promotor activity. If *ν* ≫ *λ* and *K* ≫ *δ*, the distribution of transcript abundances is the negative binomial (***Ham et al., 2019***),

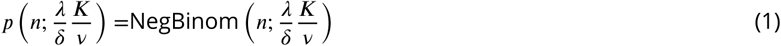

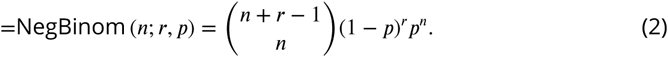

We additionally take extrinsic noise arising from variation in *glnK* (and potentially other factors) to act as a variable activation rate. This is incorporated by taking the parameter *λ* to itself vary between cells according to a log-normal distribution with mean *λ* and standard deviation *σ*. This leads to the compound distribution,

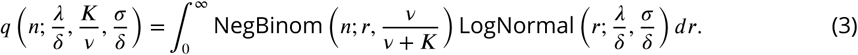

This integral can be evaluated numerically by computing the distribution for a range of values for *r* then performing a numerical integration across them. The model therefore gives us an expected transcript abundance distribution in terms of three parameter ratios, 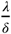 is the average normalised burst frequency, 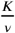 the average burst size and 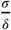 the normalised standard deviation on burst frequency. This distribution can be calculated by numerically performing the integration above.

Given this model, we fitted the parameters via a Bayesian inference approach using an MCMC sampling scheme implemented in the programming language Julia. This enabled us to obtain posterior distributions for each of the three parameter ratios, from which we obtained maximum *a posteriori* (MAP) estimates for each parameter and 95% credible intervals, as plotted in Fig. 3(C). All code relating to the modelling and parameter inference is available at https://github.com/rdbrackston/TranscriptionModels.

### Calculating extrinsic contributions to variance

Given the model fits to each dataset, we were able to calculate the extrinsic contributions to variance following the approach in ***Sherman et al.*** (***2015***); ***Hilfinger and Paulsson*** (***2011***). If *n* is the copy number of mRNA drawn from the compound distribution *q*(*n*|*λ*), where *λ* is the variable burst frequency, then the total variance of *n* may be decomposed as:

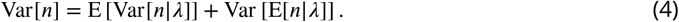

The first term is the average variance of the mRNA copy number distribution, where the average is over the distribution of values for *λ*. This term gives the contribution of intrinsic noise as it is essentially a weighted sum of the variation that arises for a fixed *λ*. The second term is the variance of the mean copy number, where the variance is again evaluated over the distribution of values for *λ*. This quantifies the contribution of the extrinsic noise since it is a measure of the variation in the copy number directly resulting from the variation in *λ*. We can calculate each of these terms numerically given a set of model parameters, thereby calculating the fractional extrinsic contribution to the total variance as,

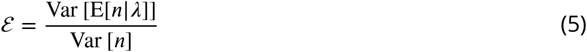

In practise the MCMC scheme yields a joint distribution over the three parameters 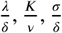. In order to accurately assess a best estimate of the extrinsic contribution as well as confidence intervals we evaluate the contribution *ε* for 4000 parameter triplets sampled from the chain. From this distribution we calculate a maximum *a posteriori* estimate and 68% credible intervals.

## Acknowledgments

This work was supported through BBSRC grants BB/N003608/1 and BB/L027135/1. TB was recipient of a Commonwealth Rutherford Fellowship. RDB is grateful to Michael Stumpf, Lucy Ham and John Pinney for many helpful discussions.

---

**Figure 1–Figure supplement 1.**
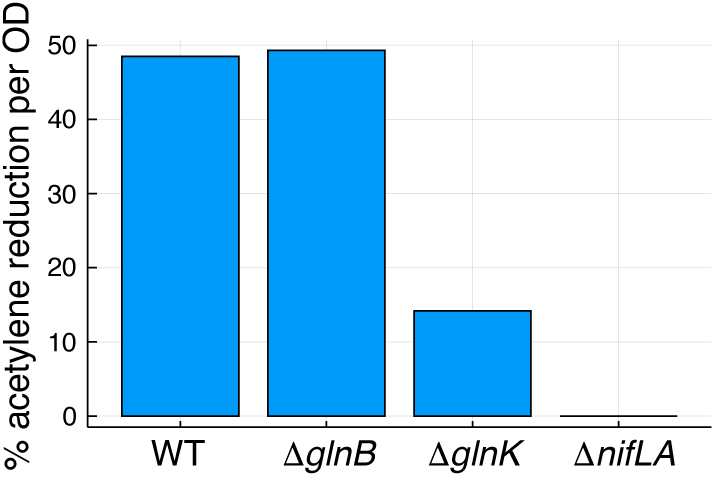
Acetylene reduction as a percentage of initial acetylene concentration, measured at 9.5 hours.

**Figure 2–Figure supplement 1.**
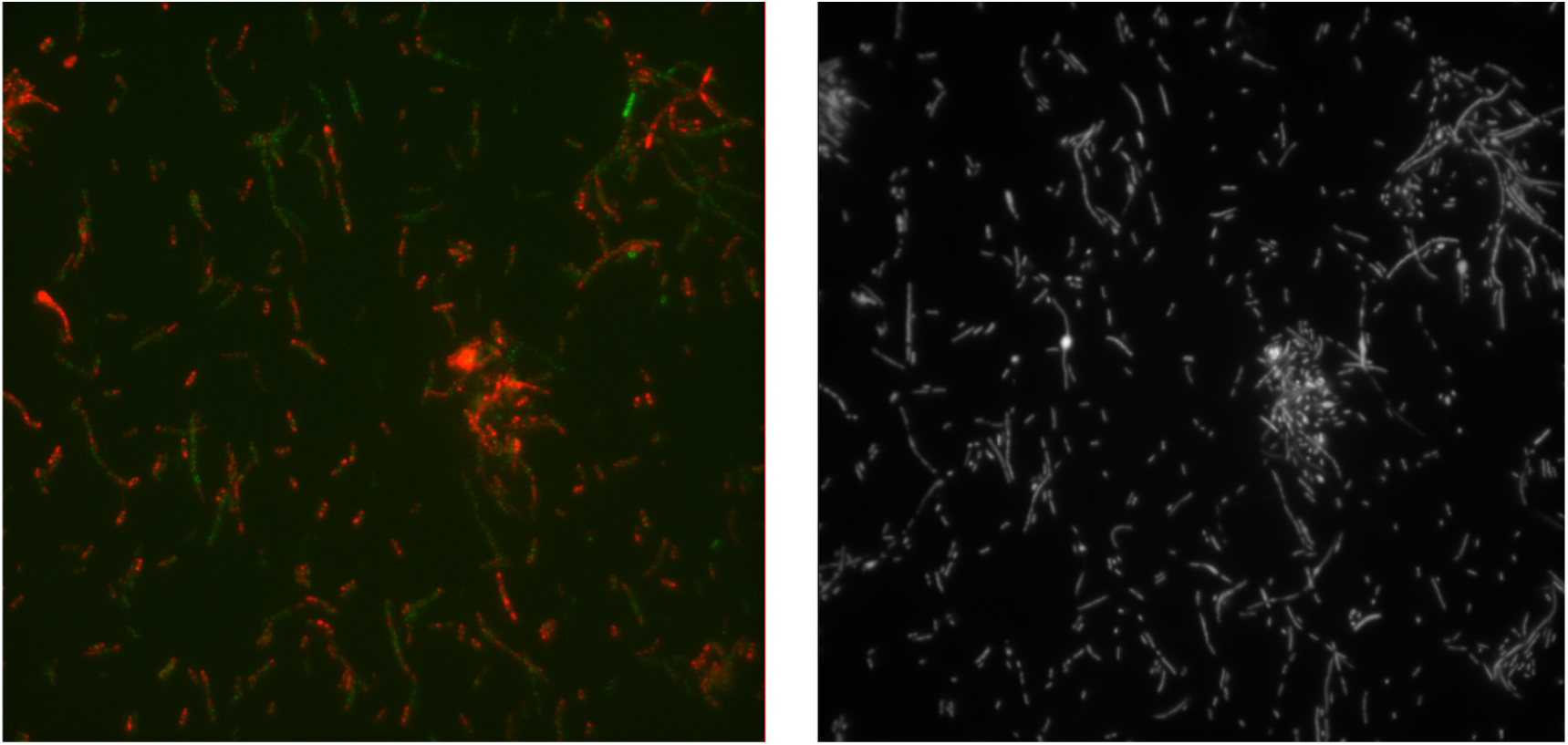
Raw images demonstrating the use of dual FISH probes. False colour luminescence from *glnK* and *nifHDK* probes (left) and DAPI stained cells (right).

**Figure 3–Figure supplement 1.**
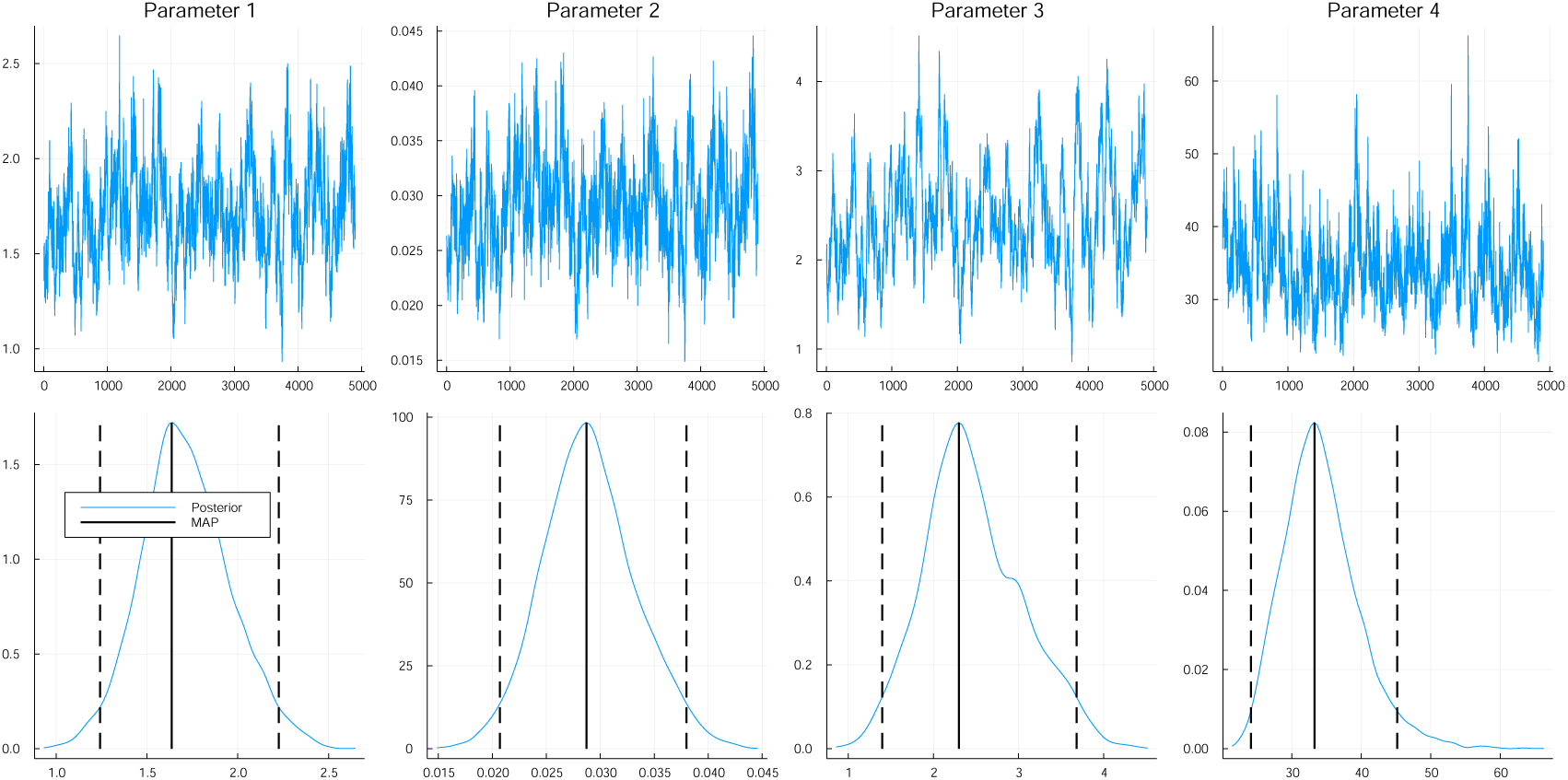
The MCMC chain is displayed in the upper figures for each of *λ*/*δ, r, σ*/*δ* and *K*/*ν*. Lower figures display the posterior distributions along with the MAP estimate and the 95% credible intervals.

